# Food-activated Microneedle Sensor for Real-time, Colorimetric Spoilage Monitoring of Pre-packaged Food

**DOI:** 10.1101/2025.07.18.665560

**Authors:** Shadman Khan, Akansha Prasad, Mahum Javed, Roderick Maclachlan, Carlos D. M. Filipe, Tohid F. Didar

## Abstract

At a time of growing food insecurity, developing technologies to reduce food waste is critical. We report an inexpensive, colorimetric spoilage sensor for real-time food product assessment. The sensor is composed of dehydrated gelatin microneedles that exhibit high mechanical integrity in their base state. However, once exposed to fluid-rich food environments, they rapidly transition to a hydrogel sensing state. Food-derived anthocyanins embedded within these microneedles enable pH-based spoilage monitoring. When applied to sealed fish products, these microneedles non-destructively penetrate through packaging and are rehydrated by the underlying fish matrix. As the product ages, a defined colour shift occurs, demonstrating strong correlation with quantitative spoilage markers. When applied to unsealed fish products for rapid testing, the large microneedle sensing interface enables accelerated colorimetric sensing. Finally, successful fresh versus spoiled categorization of smartphone-acquired images of the sensor using machine learning removes readout ambiguity, empowering consumers with independent real-time product monitoring.

## 1. Introduction

The economic and humanitarian burden of food waste has become increasingly pronounced worldwide, with an estimated one-third of edible food produced annually being discarded.^[1–4]^ Various technical measures have been pursued to reduce such waste, with sensing and eradication technologies emerging as powerful tools for the future of the food industry.^[5–10]^ Unfortunately, most developed technologies have been designed for industry-level use, leaving consumer-level waste largely unaddressed. Such waste generally originates from improper end-user food quality assessment, wherein consumers discard edible food products prematurely in the interest of food safety.^[11]^ To this end, static expiry dates are the only tool currently available to consumers to inform food edibility. These labels offer no context on the real-time state of a packaged product and come with varying definitions that lead to consumer confusion. With 20-30% of consumer-level food waste involving edible foods thrown away due to label dependency, supplying consumers with more effective tools for food quality assessment is imperative.^[11]^

From a biochemical standpoint, food spoilage is characterized by the breakdown of macromolecular textures, the overgrowth of native and contaminating microbes, and the accumulation of toxins.^[12,13]^ The complex network of molecular changes that define food spoilage makes single entity monitoring questionable. To this end, accurate spoilage tracking efforts in literature has been largely enabled through pH monitoring. Specifically, food texture degradation, microbial growth, and toxin accumulation all induce changes in pH, positioning this metric as a comprehensive measure of food quality.^[14,15]^ While food-specific pH meters are sometimes used by industry stakeholders to assess food quality, spoilage monitoring stands to offer the greatest benefit at the consumer level. Given that the use of such complex devices is not feasible on a mass market scale, developing simple, cost-effective pH sensors has been a very active area of research.^[16]^

In particular, anthocyanins – pH-responsive flavonoids derived from fruits and vegetables, have garnered significant interest.^[17]^ As these agents are native to food, they are completely food-safe and offer dramatic shifts in colour in response to single logarithmic unit changes in pH. Anthocyanins have most broadly been employed within soft, food-safe polymer films that act as entrapping matrices, without hindering the molecules’ ability to interact with fluids present in food.^[14,17,18]^ Cellulose, chitosan, and polyvinyl alcohol have been most commonly used within such systems. The non-toxic, biodegradable nature of these films ensures that they can be safely used to package foods for real-time, consumer-level monitoring. As spoilage is transduced *via* a visible colour change, consumers can assess food state without using any equipment.^[19,20]^

While intriguing, such anthocyanin-embedded spoilage sensing films offer limited real-world viability.^[21,22]^ Given that the retention of anthocyanin functionality requires these films to be highly permeable to fluids, they must be composed of porous, hydrophilic matrices. Such a composition does not sufficiently protect enclosed foods from the surrounding environment, preventing the use of such films as standalone packaging.^[14]^ Moreover, industry resistance towards increases in packaging costs makes the commercial adaptation of such technologies unlikely.

To address this, we have developed inexpensive, food-activated microneedle spoilage sensors, to be used by consumers at home (Figure 1a). The microneedle matrix is composed of gelatin – selected due to its food-safe nature, material versatility, and demonstrated applications in food sensing.^[23–25]^ Embedded with red cabbage anthocyanin, the sensors are prepared using a three-step fabrication protocol that yields dehydrated, rigid microneedles that effectively penetrate through both packaging and food. Once exposed to moisture-rich food matrices, rehydration of the gelatin matrix enables colorimetric pH monitoring (Figure 1b). These spoilage sensing performance of these sensors was evaluated using fish, given its implications in foodborne illness and well-characterized pH–spoilage correlation.^[26–28]^ The primary use-case for these sensors involves the at-home, real-time spoilage monitoring of packaged foods, wherein non-destructive insertion through overlaid packaging allows for continuous monitoring across the product’s lifespan. On-food testing yielded a gradual transition in colour from purple to blue, correlating well with changes in pH (Figure 1c-d). A second use-case involves rapid testing of opened products, wherein the significant microneedle-enabled increase in food-contacting surface area accelerates sensing. The sensors remain effective following exposure to various storage humidities and temperatures, supporting real-world use. Lastly, an artificial intelligence network was developed to automatically classify smartphone-captured images of food-embedded sensing patches, eliminating readout ambiguity and ensuring accessibility.

**Figure 1.**
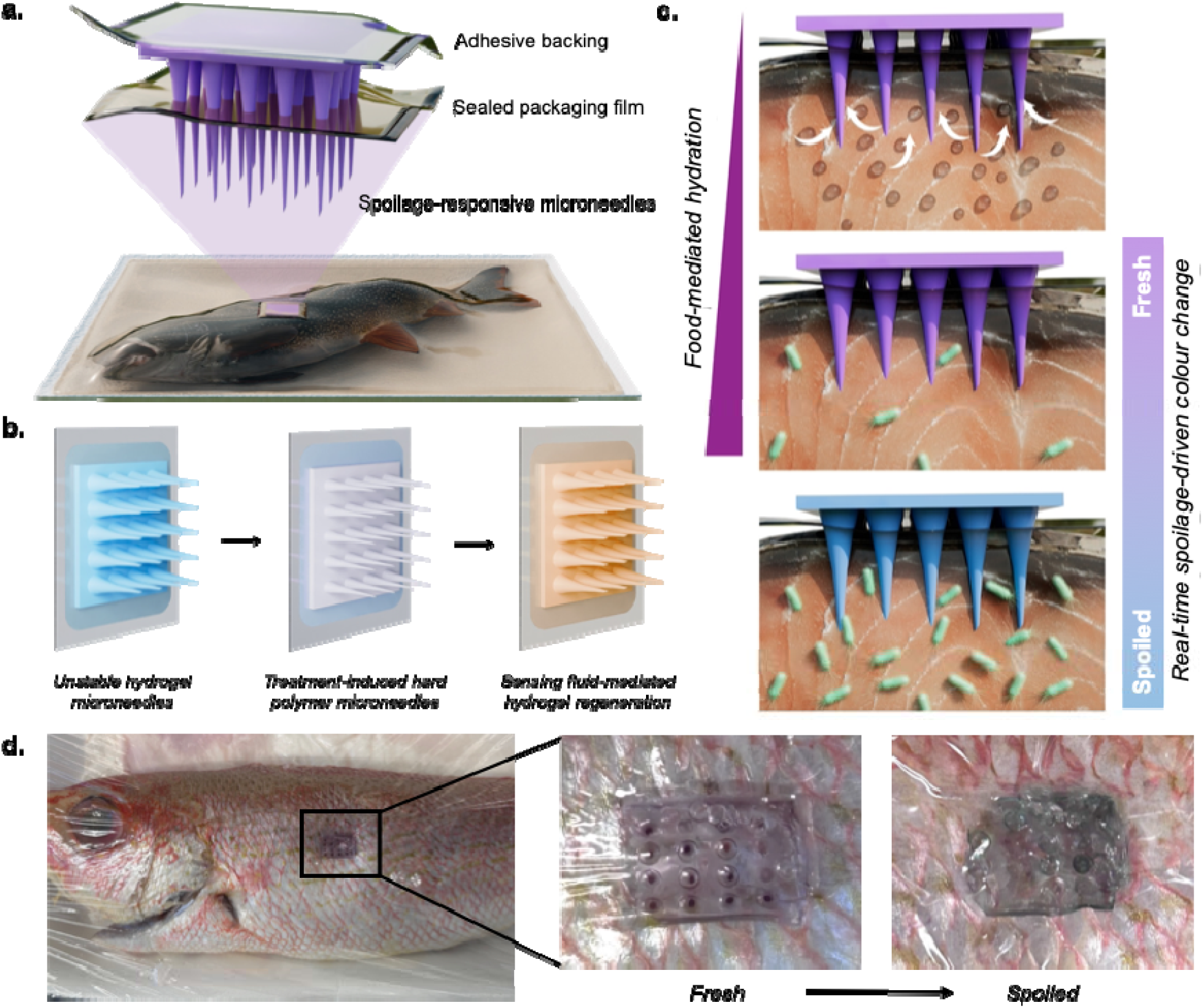
Overview of gelatin-anthocyanin fish spoilage sensor. (a) Schematic illustration of developed sensor applied to sealed fish for real-time spoilage monitoring. (b) Treatment-induced dehydration for mechanical integrity and ambient stability, followed by test fluid-induced rehydration for sensing. (c) Food-mediated hydration of the sensing microneedles, which subsequently exhibit pH-induced colour change from purple to blue as the food product spoils. (d) Proof-of-concept application showing colorimetric shift from purple to blue across fish product lifespan. Scale bar depicts 5 mm.

## 2. Results and Discussion

### 2.1 Dehydrated gelatin microneedle fabrication

Polydimethylsiloxane (PDMS) negative molds were first casted from a positive master mold with the desired dimensions, printed *via* stereolithography (Figure S1). Preliminary gelatin microneedles were fabricated through a simple casting approach (Figure 2a). Briefly, 5−20% w/v gelatin was added to deionized water, incubated at room temperature for 10 minutes to enable blooming, and then heated until completely transparent. This solution was then deposited onto the PDMS mold (Figure S2) and stored under vacuum for 30 minutes to draw out entrapped air pockets. This procedure was performed at 80^°^C to prevent premature gelation. Upon subsequent removal from heat, the casted gelatin fully solidified within 10 minutes.

**Figure 2.**
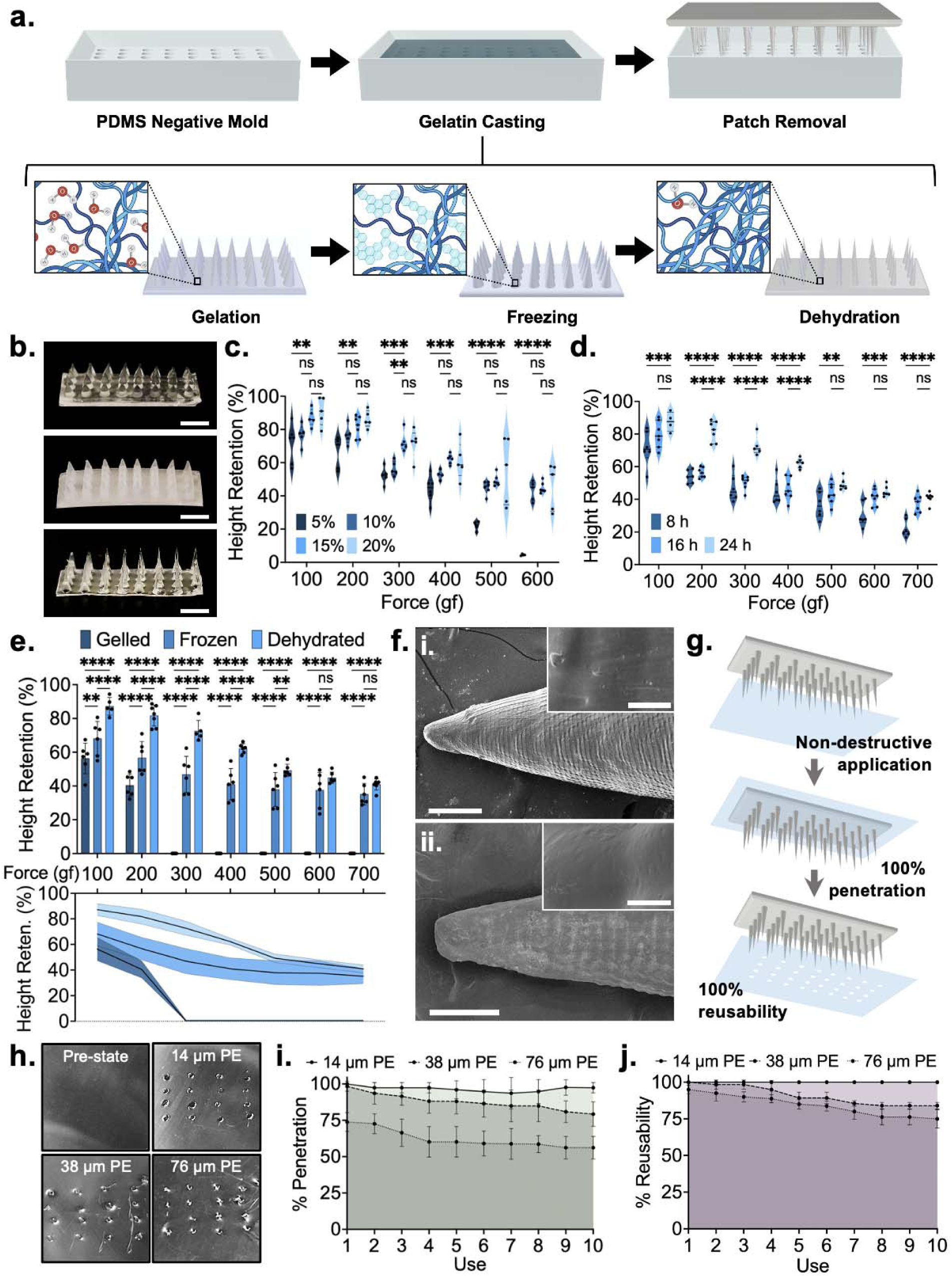
Fabrication and characterization of dehydrated gelatin microneedles. (a) Schematic illustration of fabrication protocol. (b) Optical images of gelled, frozen and dehydrated gelatin microneedles. Scale bar depicts 4 mm. (c) Optimization of gelatin concentration based on the mechanical strength of resultant microneedles. (d) Optimization of dehydration time based on the mechanical strength of resultant microneedles. (e) Baseline mechanical characterization of optimized gelled, frozen, and dehydrated microneedles. (f) Scanning electron microscopy images of (i) gelled and (ii) dehydrated microneedles. Main image scale bar depicts 150 μm, while inset image scale bar depicts 10 μm. (g) Schematic illustration of film penetration testing. (h) Optical images microneedles of food packaging films penetrated by dehydrated gelatin microneedles. Scale bar depicts 3 mm. (i) Percent penetration of dehydrated gelatin microneedles through food packaging films through ten uses. (j) Percent reusability of dehydrated gelatin microneedles through food packaging films through ten uses. All error bars depict standard deviation. All asterisks represent significant differences at corresponding significance levels.

Reversible dehydration was enacted through treatments that succeeded gelation. Specifically, the casted gelatin microneedles were first stored at −20^°^C overnight to induce freezing. The needles were then incubated at 30^°^C overnight, yielding the final dehydrated microneedle patch (Figure 2b). This freeze-thaw approach is necessary to induce microstructural changes that increase microneedle strength, fluid absorption capacity, and dehydration efficacy. Specifically, as the water within the gelatin matrix freezes, it forms large, disorganized ice grains.^[29]^ These grains condense surrounding polymer chains, facilitating physical crosslinking and the formation of inter-macromolecular bonds. This in turn, increases the microneedles’ mechanical integrity. Moreover, the cavities produced by these large ice grains do not dissipate following thawing of the ice grains,^[29]^ resulting in increased liquid absorption capacity relative to non-frozen gelatin microneedles. Finally, water molecules that are entrapped within the core of individual ice grains are unable to form hydrogen bonds with surrounding gelatin chains.^[29]^ These water molecules are thus largely unstable within the polymer matrix, making them prone to dehydration. Importantly, the microneedle patches were only removed from the PDMS negative mold following dehydration, wherein the mold acted as a scaffold for the microneedles during freezing and dehydration.

Mechanical optimization of the dehydrated microneedles involved considerations towards gelatin concentration in the precursor solution and dehydration time (Table S1). Sample conditions assessed through the microneedle height reduction observed following the application of perpendicular mechanical force, wherein a lower height reduction signals higher mechanical integrity. Gelatin w/v concentrations of 5%, 10%, 15%, and 20% were evaluated at forces ranging from 100 to 600 gram-force (gf) (Figure 2c). Dehydrated 5% gelatin microneedles exhibited significantly reduced mechanical integrity across all tested force conditions (*P* < 0.01, 0.01, 0.001, 0.001, 0.0001, 0.0001) compared to dehydrated 15% microneedles. Between the dehydrated 10%, 15% and 20% microneedles, a slight decrease in mean height reduction was observed as concentration was increased. While these differences were generally not statistically significant, 10% dehydrated microneedles were noted to be slightly inferior to 15% and 20% dehydrates microneedles due to their higher height reduction mean values. While 15% and 20% dehydrates microneedles were largely similar with regards to their mean values, the latter exhibited higher variance in the degree of height reduction. It was noted that 20% gelatin concentrations yielded increasingly brittle properties, making the needles prone to snapping upon force application. As such, the 15% gelatin concentration was selected, wherein a height reduction of 12.77% ± 5.20 was observed at 100 gf and a height reduction of 55.10% ± 3.33 was observed at 600 gf.

To assess the effect of dehydration time on mechanical strength, samples dehydrated for 8 hours, 16 hours, and 24 hours were evaluated at forces between 100 and 700 gf (Figure 2d). Compared to the 24-hour samples, 8-hour samples exhibited significantly higher height reduction at all tested forces (*P* < 0.001, 0.0001, 0.0001, 0.0001, 0.01, 0.001, 0.0001). Samples dehydrated for 16 hours demonstrated a significant improvement in mechanical strength relative to the 8-hour samples but were still significantly weaker than 24-hour samples at forces between 200 and 400 gf (*P* < 0.0001, 0.0001, 0.0001). While differences were not significant at the other tested forces, the 24-hour samples consistently exhibited a mean height reduction lower than the 16-hour samples. Improvements in strength were not observed when dehydration was performed for longer than 24 hours. As such, 24 hours was noted to be the optimal dehydration time, wherein height reductions of 12.77% ± 5.20 and 59.06% ± 3.05 were observed at 100 gf and 700 gf, respectively.

With optimized fabrication parameters selected, an assessment of microneedle integrity was performed as each stage of post-gelation processing to understand the impact of freezing and dehydration (Figure 2e). Gelled microneedles offered very low mechanical strength, with height reductions of 43.64% ± 9.03 and 55.55% ± 6.65 at forces of 100 gf and 200 gf, respectively. This was significantly higher than both their frozen (31.80% ± 10.05, *P* < 0.01; 43.28% ± 9.61, *P* < 0.0001) and dehydrated counterparts (12.77% ± 5.20, *P* < 0.0001; 18.25% ± 6.24, *P* < 0.0001). Complete height reduction was observed at forces of 300 gf and above. Conversely, frozen microneedles demonstrated rapidly deteriorating mechanical integrity after removal from their −20^°^C treatment environment. Their exterior surface thawed almost immediately, while the inner core thawed within two minutes. Accordingly, these microneedles exhibited significantly higher height reductions at forces between 100 and 500 gf compared to dehydrated microneedles (*P* < 0.0001, 0.0001, 0.0001, 0.0001, 0.01), as the thawed exterior offered little structural resistance to applied forces. At forces of 600 and 700 gf, the difference between the two conditions was not significant, owing to the temporarily intact ice core. Still, the mean height reduction values exhibited by the dehydrated microneedles at these forces were still lower than that of frozen microneedles. The dehydrated microneedles thus presented a breakthrough material that offered mechanical integrity superior to that of frozen microneedles, in a format that remains stable indefinitely at room temperature.

The gelled and dehydrated microneedles were subjected to scanning electron microscopy (SEM) (Figure 2f). Dehydration was signified by a significant reduction in microneedle diameter, alongside microscale textures that was attributed to gelatin crystallization. When tested against target matrices, the dehydrated microneedles exhibited excellent penetration of polyethylene food packaging films with thicknesses of 14μm, 38μm and 76μm (Figure 2g-h). Effective penetration was also observed with alternative packaging materials and non-target food products (Figure S3-4). The packaging penetration capabilities of the dehydrated microneedles was quantified through the percent of applied microneedles that effectively penetrated through the three polyethylene films (Figure 2i). Single patches were tested over ten times to simultaneously assess the microneedles’ resilience, with the number of reusable needles – total needles minus broken needles, noted after each trial (Figure 2j). With 14μm thick polyethylene, the penetration rate was consistently above 93% and reusability was noted to be 100.0% following ten trials. 38μm thick polyethylene films offered slightly more resistance, yielding 79.3% penetration by the tenth trial, and an 83.7% reusability rate. Finally, testing with 76μm films resulted in a 56.2% penetration rate and a 75% reusability rate. Importantly, rates were calculated with reference to the number of microneedles at the start of the first use. When adjusted to the number of intact needles at the start of the tenth use, the penetration rates for 38μm and 76μm films are 94.7% and 74.9%, respectively. While the use of these microneedles for real-time spoilage sensing would only require a single penetration event, these studies demonstrate the mechanical robustness of the developed structures.

### 2.2 Anthocyanin integration and optimization

Anthocyanin integration within the dehydrated gelatin microneedle matrix represented the next objective. Red cabbage anthocyanin was suitable due to its defined colour shift from purple to blue at pH 7.0, which corresponds with the fish spoilage threshold reported in prior literature.^[30,31]^ Anthocyanin powder was first dissolved within deionized water to create a 1% w/v stock solution. This solution was then further diluted to 0.1%, 0.5%, and 0.9% in deionized water. Here, it was hypothesized that too low of a concentration would offer a suboptimal colour shift, while too high of a concentration would buffer the colour shift beyond the desired spoilage threshold. The change in absorbance exhibited by these three anthocyanin concentrations when exposed to fresh versus spoiled fish was quantified (Figure 3a). Measurements were taken at several wavelengths to identify the optimal parameters for colour shift monitoring, wherein absorbance measured at 300 nm and 350 nm exhibited the most dramatic changes. While the differences between conditions were statistically insignificant, increased anthocyanin concentrations yielded slightly higher mean shifts in absorbance at 300 nm and 350 nm. Next, we sought to assess how shifts in absorbance correlated with pH on a day-to-day basis throughout the products’ lifespan. Regression analyses were performed between the absorbance of anthocyanin solutions incubated with fish for 30 minutes and fish pH, over the course of 6 days (Figure 3b). Here, 0.1% anthocyanin solutions exhibited the lowest correlation with fish pH, with r-values of 0.8557 and 0.8424 at wavelengths of 300 nm and 350 nm, respectively (*P* < 0.05, 0.05). On the other hand, the changes in absorbance exhibited by 0.5% and 0.9% anthocyanin solutions demonstrated strong correlation with fish pH, with r-values all above 0.95 (*P* < 0.01, 0.01, 0.01). Importantly, 0.9% anthocyanin solutions were only quantified at 350 nm, as overflow was noted at 300 nm.

**Figure 3.**
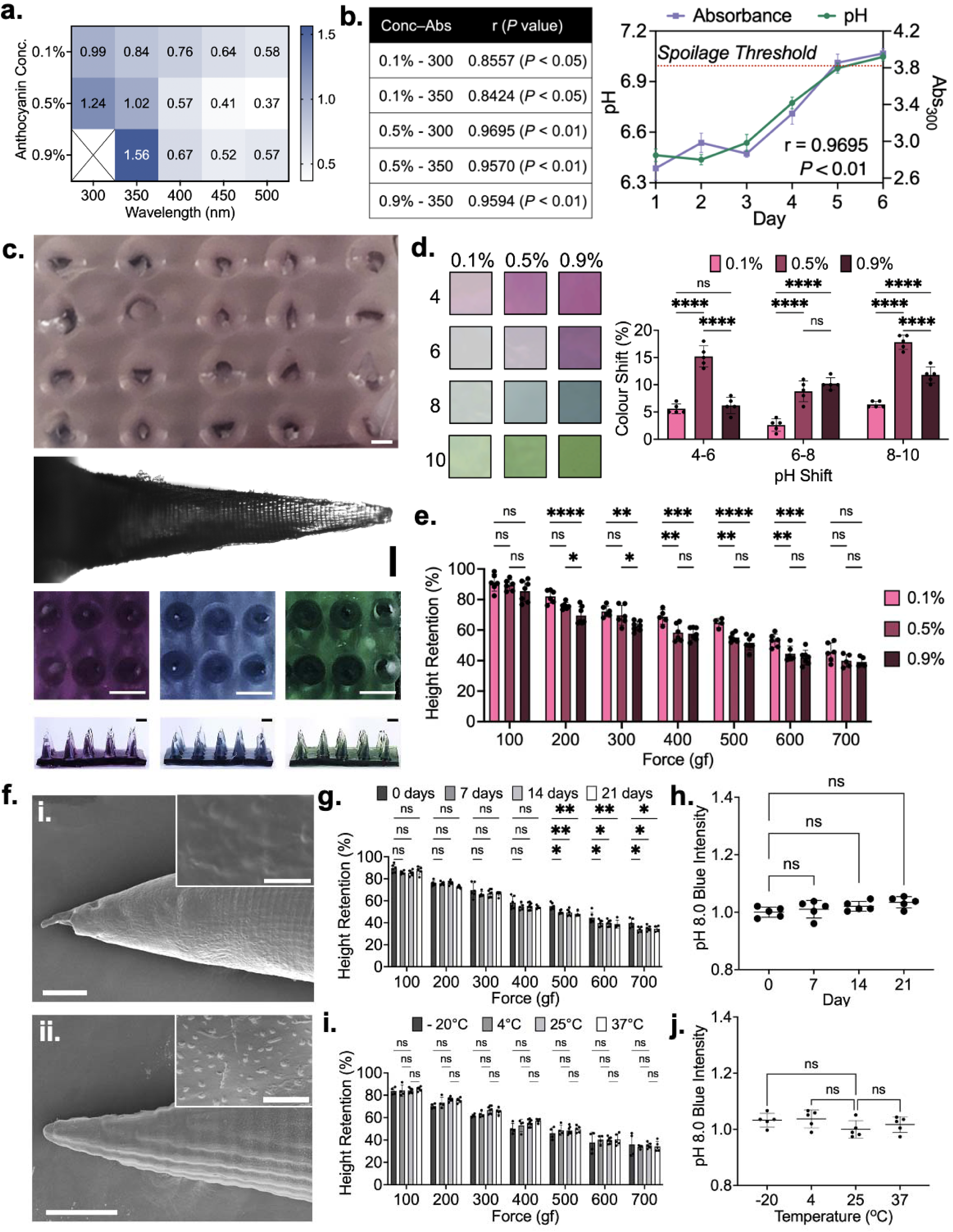
Anthocyanin incorporation into dehydrated gelatin microneedles and corresponding performance testing. (a) Absorbance shifts observed through fish spoilage using various anthocyanin concentrations and measurement wavelengths. (b) Correlation analysis of anthocyanin concentration-absorbance wavelength combinations. (c) Optical images of an anthocyanin-embedded dehydrated gelatin microneedle sensor, a single needle, and rehydrated patches following exposure to pH. 6.0, 8.0, and 10.0, from left to right. Coloured images have scale bars depicting 1 mm, while Brightfield image scale bar depicts 200 μm. (d) Percent colour shift exhibited by sensors prepared using varying anthocyanin concentrations at different pH ranges. (e) Mechanical assessment of anthocyanin-embedded microneedle sensor. (f) Scanning electron microscopy images of (i) gelled and (ii) dehydrated anthocyanin-embedded microneedles. Main image scale bar depicts 150 μm, while inset image scale bar depicts 10 μm. (g) Mechanical integrity of sensors after long-term storage. (h) pH responsiveness of sensors after long-term storage. (i) Mechanical integrity of sensors after storage at diverse temperature for 5 days. (j) pH responsiveness of sensors after storage at diverse temperature for 5 days. All error bars depict standard deviation. All asterisks represent statistically significant differences at corresponding significance levels.

Recognizing that the dehydrated gelatin matrix may alter the colour-responsive properties of embedded anthocyanin, subsequent studies employed dehydrated gelatin microneedles embedded with the pH-responsive agent. In its initial state, the patch exhibits a faint purple hue (Figure 3c). However, upon incubation within solutions with pH values of 6.0, 8.0, and 10.0, expected shifts to dark purple, blue, and green were observed in under five minutes. Importantly, this was the first indication that anthocyanin embedded within dehydrated gelatin remained fully functional and accessible for pH sensing. In fact, it was observed that the microneedles returned to a largely rehydrated state, similar to their initial gelled state. To this end, the rate of rehydration was evaluated on both short and long-term timescales (Figure S5-6).

To quantify the pH responsiveness of anthocyanins embedded within dehydrated gelatin matrices, the percent colour shift was calculated between samples incubated at pH values of 4.0, 6.0, 8.0, and 10.0 (Figure 3d). When pH increased from 4 to 6 and from 8 to 10, the shifts exhibited by the 0.5% dehydrated anthocyanin patches were significantly more dramatic than that of 0.1% (*P* < 0.0001, 0.0001) and 0.9% patches (*P* < 0.0001, 0.0001). With regards to the pH 6.0 to 8.0 transition – the range most relevant to fish spoilage, the 0.5% dehydrated patches demonstrated a much more substantial 8.8% ± 1.7 mean shift in colour compared to the 2.6% ± 1.3 mean shift exhibited by 0.1% patches (*P* < 0.0001). The 10.2% ± 1.3 mean shift exhibited by the 0.9% patches was not significantly different from the 0.5% patches, making both relevant for application-specific testing.

### 2.3 Material assessment of spoilage-sensing microneedles

To ensure that the introduction of anthocyanin at high concentrations does not yield significant alterations to the mechanical integrity of dehydrated gelatin microneedles, force testing was performed on dehydrated microneedles embedded with 0.1%, 0.5%, and 0.9% anthocyanin (Figure 3e). While samples embedded with higher anthocyanin concentrations exhibited slightly higher microneedle height reductions in response to the application of a perpendicular force, the changes were small in magnitude. Specifically, 0.1% and 0.9% dehydrated anthocyanin microneedles differed by less than 14% across all tested conditions.

Interestingly, 0.1% dehydrated anthocyanin microneedles generally exhibited less height reduction than dehydrated anthocyanin-free microneedles. This improvement in mechanical integrity is attributed to increased structural stability afforded by hydrogen bonding between free anthocyanins and the polymeric microneedle matrix. While this would suggest that 0.5% and 0.9% dehydrated anthocyanin microneedles would exhibit even greater mechanical integrity, it is hypothesized that the abundant free anthocyanins present within these samples engage in interactions with water molecules present within the gelled and frozen microneedle states. Bonded water molecules are resistant to extraction *via* dehydration, yielding a dehydrated material with slightly higher moisture content. Owing to these counteracting effects, there is little difference in the mechanical integrity of dehydrated anthocyanin-free and dehydrated 0.9% anthocyanin microneedles. Specifically, the two respective microneedle conditions demonstrate height reductions of 12.77% ± 5.20 and 14.51% ± 6.94 at 100 gf, alongside reductions of 59.06% ± 3.05 and 60.87% ± 3.13 at 700 gf. The hypothesized increase in anthocyanin-driven intermolecular interactions is supported by SEM images of anthocyanin-embedded microneedles (Figure 3f). Dehydrated microneedles with embedded with anthocyanins exhibit a significant increase in surface textures attributed to gelatin crystallization.

Next, the impact of long-term storage on the mechanical integrity and pH responsiveness of dehydrated, anthocyanin-embedded microneedles was assessed. Mechanical integrity remained largely unchanged for 0.5% anthocyanin-embedded microneedles, with no significant differences observed amongst all storage samples below 400 gf (Figure 3g). While slight increases in mean height reduction were observed with long-term storage between forces of 500 and 700 gf (*P* < 0.01−0.05), these differences did not exceed 8% under any tested conditions. With regards to on-package application, such a difference would induce negligible changes in penetration and reusability rates. With regards to pH-responsiveness, 0.5% anthocyanin-embedded sensing patches were stored for 7, 14, and 21 days in ambient conditions. These samples were subsequently incubated in pH 8.0 standard solution for 30 minutes, with the resultant blue colour intensity quantified (Figure 3h). No significant differences in the blue colour output were observed, confirming that the dehydrated gelatin environment did not influence anthocyanin functionality. As the maintenance of an effective colorimetric response was confirmed in the lower 0.5% concentration condition, only the mechanical integrity of 0.9% anthocyanin-embedded microneedles was assessed. These samples also exhibited little degradation in mechanical integrity over time (Figure S7).

Finally, the impact of temperature on the mechanical integrity and pH responsiveness of dehydrated, anthocyanin-embedded microneedles was evaluated. 0.5% anthocyanin-embedded sensing patches were stored at −20^°^C, 4^°^C, 25^°^C, and 37^°^C for 7 days and subsequently tested using the same approaches used to assess the long-term storage samples. Mechanical stability did not significantly differ across any of the tested conditions and no significant changes in blue colour output were observed (Figure 3i-j). The impacts of higher temperature conditions and varying humidities were also assessed, where no degradation in performance was observed (Figure S8-9). Force testing of 0.9% anthocyanin-embedded microneedles following variable temperature storage also yielded largely insignificant changes to mechanical integrity (Figure S10).

### 2.4 Proof-of-concept testing and machine learning integration

Next, we sought to investigate the real-time fish spoilage sensing properties of the developed microneedles, wherein the two aforementioned consumer-centric use cases were considered. With regards to non-destructive penetration of sealed food packaging for at-home, real-time spoilage monitoring, the 0.5% anthocyanin-embedded spoilage sensing patches behaved as predicted (Figure 4a). When packaged fish samples remained edible (Days 1 and 2), optical images collected using a smartphone depict patches with a defined purple hue. However, once the spoilage threshold of 7.0 was passed on Day 3, a dramatic shift to blue was observed, which became increasingly pronounced on Day 4. A standard RGB scale was employed to quantify the loss of red colour intensity, as a function of the overall spoiled-induced colour shift exhibited by the microneedle spoilage sensors. Red colour intensity averaged arbitrary unit values of 99.6 ± 3.4, 87.1 ± 4.2, 66.6 ± 4.9, and 57.1 ± 6.0 on Days 1 through 4, respectively. Contrarily, 0.9% anthocyanin-embedded spoilage sensing patches did not offer effective performance. This is attributed to the fluid content available within the fish matrix being insufficient for the induction of colour change within the highly concentrated anthocyanin environment present within these samples.

**Figure 4.**
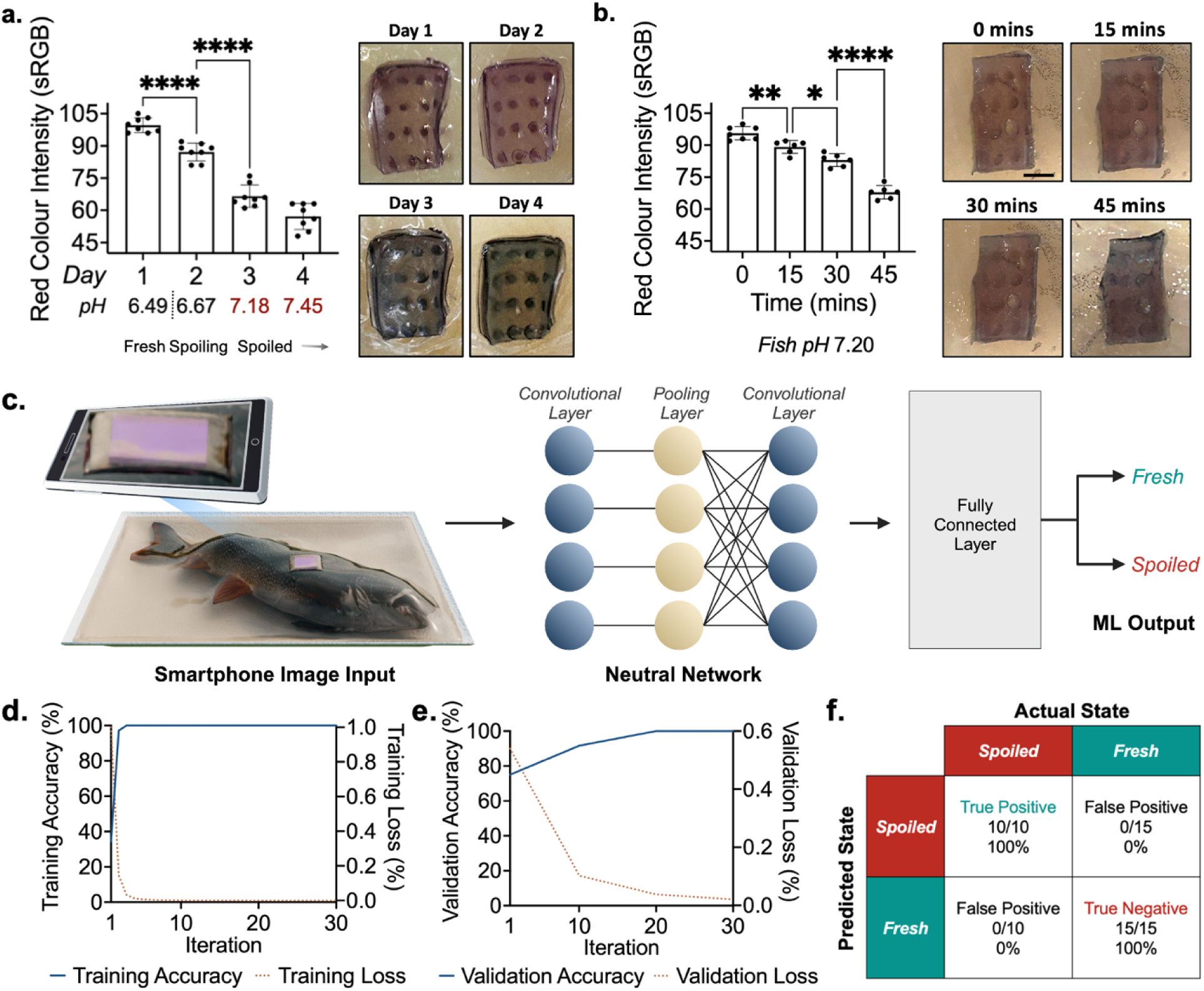
Proof-of-concept testing and machine learning integration. (a) Quantification and associated images of real-time monitoring of sealed fish throughout its product lifespan. (b) Quantification and associated images of spoiled fish rapid test. (c) Schematic illustration of real-world, consumer-level use, yielding a fresh or spoiled readout. (d) Training accuracy and loss curves. (e) Validation accuracy and loss curves. (f) Confusion matrix of AI model when tested with 25 smartphone-captured images.

With regards to an at-home rapid test for opened products, spoiled fish samples were penetrated with the spoilage sensors and imaged with a smartphone every 15 minutes (Figure 4b). Again, a pronounced shift from purple to blue was observed. The mean red colour intensity was quantified according to a standard RGB scale, yielding arbitrary unit values of 96.0 ± 3.1, 89.2 ± 3.1, 84.8 ± 3.2, and 67.8 ± 3.2 at 0, 15, 30, and 45 minutes, respectively. Significant shifts were observed between each respective timepoint (*P* < 0.01, 0.05, 0.0001). Recognizing that passing the spoilage threshold in the day-scale monitoring study corresponded with a red colour intensity of 66.6 ± 4.9, 45 minutes was noted to be the required time for rapid test sensing. That being said, the significant downward trend exhibited at the 15 and 30-minute marks suggests that rate-change monitoring may enable spoilage assessment within a shorter time span. Again, 0.9% anthocyanin-embedded spoilage sensing patches were not effective. Importantly, a colorimetric response was not elicited by flat, dehydrated gelatin patches within this sensing timeframe, showcasing the importance of microneedle-enabled surface area enhancement for rapid testing (Figure S11).

To automate our solution and ensure it was user-intuitive and accessible for users with limited vision, a convolutional neural network (CNN)-based artificial intelligence model was trained using smartphone-acquired microneedle images to provide a simple “Fresh” or “Spoiled” classification based on a given microneedle patch image (Figure 4c). A CNN was chosen given its suitability with image-based input and classification.^[32]^ Images were first randomly split into training and validation datasets and then used to train the model across 30 iterations. Validation was performed on every 10th iteration to ensure adequate training and model improvements.

Overall, the trained model was highly accurate (Figure 4d) and importantly, this high accuracy was maintained across validation (Figure 4e). Similarly, training and validation loss both approached zero, indicative of the model’s precision. To further test the broad applicability of th model, blind testing was performed to a separate dataset from the training and validation ones. This testing yielded true positive and true negative rates of 100% across all 25 images that were inputted, with no false negative or false positive classifications (Figure 4f). Such an AI model simplifies sensor readout by using a simple smartphone-acquired optical image, thus eliminating uncertainty in colorimetric interpretation and making the sensor more accessible and user-friendly.

## 3. Conclusion

In this work, we introduce pure gelatin microneedles that offer shifts in their mechanical properties in response to environmental stimuli. The microneedles are produced through a three-step process that involves gelation, freezing, and dehydration. In their dehydrated state, the microneedles exhibit robust mechanical integrity – similar to that of frozen needles, but in a shelf-stable form. Their excellent physical properties allow them to effectively penetrate through food packaging in a non-destructive fashion. Only when they contact fluids, do these dehydrated gelatin microneedles return to a hydrogel state more typical of gelatin. When embedded with pH-responsive, colour-changing anthocyanins, this rehydration event makes anthocyanins accessible to the hydrating fluid, permitting colorimetric sensing. Long-term storage and food-relevant temperature exposure both yielded limited changes to both the mechanical integrity and pH responsiveness of the developed sensor. When applied towards real-time monitoring of packaged fish, the developed spoilage sensor exhibited robust correlation with spoilage events, offering dramatic shifts from purple to blue. In another vein, when applied directly onto spoiled fish as a rapid test to be used prior to consumption, a complete shift to blue is observed within 45 minutes. Finally, the integration of a machine learning model enabled spoilage assessment using a smartphone, preventing readout ambiguity and increasing accessibility. Looking forward, while not obvious to the naked eye, subtle shifts in colour occur prior a full transition to blue. This suggests that sensor performance may be readily improved through further training of the artificial intelligence model. In the case of real-time packaged food monitoring, this may enact a “spoiling” classification to convey to the consumer that a product is approaching spoilage. With regards to rapid testing, such efforts stand to reduce detection time. Yet even in its current state, the developed sensor stands to offer consumers with the power to effectively and accessibly assess the quality of purchased foods.

## 4. Methods

### Materials

Polydimethylsiloxane (SYLGARD 184) was purchased from Dow Corning (Michigan, United States). Gelatin was purchased from Fortino’s (Ontario, Canada). Polyethylene wraps were sourced from Thomas Scientific (New Jersey, United States) and Walmart Canada (Ontario, Canada). pH standard solutions were sourced from Millipore Sigma (Ontario, Canada). Haddock fillet samples were purchased from Food Basics (Ontario, Canada). *Resin master mold fabrication.* A CAD model of a microneedle mold was designed in Autodesk Inventor, ensuring precise dimensions and features. The 3D model file was then converted into an STL file, recognized by the stereolithography (SLA) 3D printer. Black resin was used as the printing material, and the printer created the mold layer by layer, resulting in a detailed and accurate microneedle mold. Once the print was complete, it was rinsed with 100% isopropyl alcohol to remove excess resin on the surface and then dried. Post-curing of the molds involved placing the printed parts in a water bath and exposing them to UV light on both sides for 10 minutes, followed by overnight drying at 30. This post-curing process maximized the mechanical strength and ensured stability of the molds.

### PDMS negative mold fabrication

PDMS Sylgard 184 was prepared at a 10:1 ratio of base resin to curing agent, stirred for 10 mins, and then desiccated for 30 mins. This mixture was then casted onto the resin master molds and desiccated again for 30 mins. The casted PDMS were then cured at 60^°^C overnight and then detached from the master mold to obtain the PDMS negative mold.

### Gelled microneedle fabrication, freezing, and dehydration

Gelatin powder was added to deionized water at 5%, 10%, 15%, and 20% w/v concentrations. The mixtures were then stirred and left to bloom for 10 mins and then heated until transparent using a conventional microwave. The resultant solution was casted onto pre-heated PDMS negative molds and placed under vacuum at −0.08 MPa for 5 mins at 80. Any entrapped air pockets visible after desiccation were physically agitated with a needle and removed. The casted solutions were left to gel for 10 minutes before proceeding to post-fabrication treatment steps. At this point, the gelled microneedles were left adhered to the negative mold and placed into a conventional freezer set to −20 for 24 h. The microneedles-mold pairing was then subsequently placed onto a hot plate set to 30 for 8 h, 16 h, or 24 h dehydration times. Microneedles were separated from the mold only after dehydration. Anthocyanin-embedded microneedle fabrication followed the same protocol, but used 0.1%, 0.5%, and 0.9% anthocyanin solutions instead of deionized water.

### Microneedle imaging

Colour optical images of the microneedle patches were taken using a Samsung Galaxy S21 Ultra smartphone. Brightfield imaging of a single needle was performed using a Nikon Eclipse Ti2 inverted microscope.

### Mechanical testing

In preparation for mechanical testing, each microneedle patch was precisely cut into rows of four needles. To evaluate mechanical integrity, the force was normalized to range from 100 gf to 700 gf per individual needle. High-resolution images of the needles were taken before any force application to document the baseline structure. A Biomomentum Mach 1 mechanical tester was used to apply perpendicular incremental forces, ranging from 100 gf to 700 gf, to the microneedles. After each subsequent increase in force, additional images were captured to analyze any deformations or structural failures. Using these images, the height reduction of the individual microneedles was then measured to assess mechanical strength. Lower height reductions indicated higher mechanical integrity. Data from different force levels were compared to evaluate the mechanical properties of the microneedles under varying stress conditions. This standardized method ensured consistent evaluation of microneedle mechanical properties across multiple samples with varying conditions.

### Scanning electron microscopy

Single microneedles were isolated from their base to induce a two-dimensional form factor that was better suited for top-down perspective imaging. Samples were mounted using carbon tape and nickel paste. A sputter coater (Polaron E1500, Polaron Equipment Ltd., Watford, Hertfordshire) was subsequently used to coat the samples with 5 nm of platinum. Samples were imaged using the JEOL JSM-7000F.

### Penetration and reusability studies

Polyethylene films of varying thickness were secured across the top of a cylindrical support. Microneedle patches were then placed on top of the films with the needles facing down. A mechanical tester was used to apply a consistent force of 500 gf. The patches were then removed, and the number of penetration sites was quantified to determine the penetration rate. This number of undamaged individual microneedles in a patch was also quantified to determine the patch reusability rate. Both rates were calculated as a function of the initial needle count. Patches were subjected to 10 successive trials each.

### Anthocyanin absorbance versus pH studies

Haddock fish fillets purchased from a local grocery store were divided into 5 g samples and individually stored in sealed bags. Three samples were subjected to pH assessment each day using a Mettler Toledo SevenExcellence pH meter. Next, 0.5 g portions were isolated from each sample for anthocyanin colour change assessment. These portions were added to 1 mL aliquots of anthocyanin solutions with concentrations of 0.1%, 0.5%, and 0.9% and incubated for 30 mins. 100μL samples of the tested solutions were then added to a well plate for absorbance measurements at various wavelengths. Absorbance readings were taken using a Synergy Neo2 plate reader.

### Colour shift studies on dehydrated anthocyanin-embedded patches

Dehydrated anthocyanin-embedded patches were subjected to 4.0, 6.0, 8.0, and 10.0 pH standard solutions. After 15 mins of incubation, the samples were imaged using an Epson Perfection V850 Pro Scanner. Quantification of colour shifts involved viewing each colour as a three-dimensional coordinate on an RGB colour map. From here, the distance between two colours was quantified as:

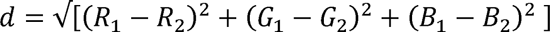

The distance was then referenced as a percent of the maximum distance between two colours:

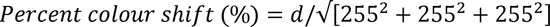

### Day-scale monitoring of sealed fish products

Haddock fish fillets purchased from a local grocery store were divided into 5 g samples and individually wrapped with polyethylene packaging film. Microneedle spoilage sensors were inserted through the packaging film, into the underlying fish matrix. A secondary film was secured overtop the sensor to prevent unwanted air exposure. The samples were stored in the fridge and imaged using a smartphone every day, until spoilage was well-established. Quantification of colour change involved assessing the intensity of red colour at the microneedle dots observable through the underside of the patch. As the fish spoiled, a loss in red colour intensity was observed.

### Minute-scale monitoring of opened fish products

Haddock fish fillets purchased from a local grocery store were divided into 5 g samples and stored until spoiled. The samples were then penetrated with microneedle spoilage sensors and imaged every 15 minutes. Again, quantification of colour change involved assessing the intensity of red colour at the microneedle dots observable through the patch, wherein a loss in intensity was observed as the microneedle patches rehydrated.

### Storage and temperature stability studies

Microneedle spoilage sensors were stored under ambient conditions for 7, 14, and 21 days. They were then subjected to mechanical testing, as described above. pH responsiveness was assessed through 20 mins incubation in pH 8.0 solution, after which samples were scanned using an Epson Perfection V850 Pro scanner. Quantification involved assessing the blue colour intensity at the microneedle dot sites observable through the patch.

### CNN development

The CNN model was trained and developed using MATLAB’s MathWorks Deep Learning using transfer learning from a pre-trained deep CNN. A training dataset of 30 images were used with a 60/40 training to validation split. All images were acquired using a Apple iPhone 14 smartphone and cropped to remove unnecessary background. 30 training iterations were used with validation performed on every 10th iteration.

## Acknowledgements

S.K. and A.P. are recipients of the Vanier Canada Graduate Scholarship funded by the Natural Sciences and Engineering Research Council (NSERC). This research was undertaken, in part, thanks to the Canada Research Chairs Program awards to T.F. Didar. T.F. Didar is funded by the Natural Sciences and Engineering Research Council (NSERC) through the Canada Discovery Grant and the Ontario Early Researcher Award. This project also received funding from Toyota Tsusho Canada Incorporated awarded to T.F.D.

## Supporting Information

Supporting Information is available from the author.

