## Supplementary Information for "Food-activated Microneedle Sensor for Real-time, Colorimetric Spoilage Monitoring of Pre-packaged Food"

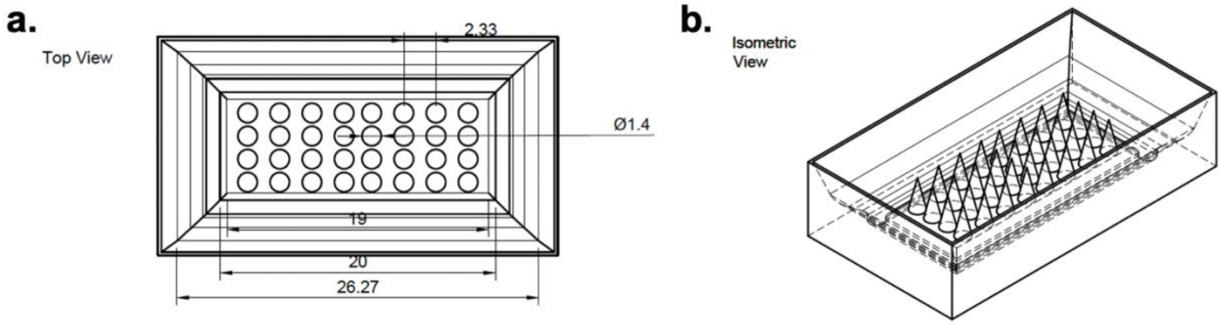


**Figure S1.** Schematic illustrations of master mold fabricated *via* stereolithography.

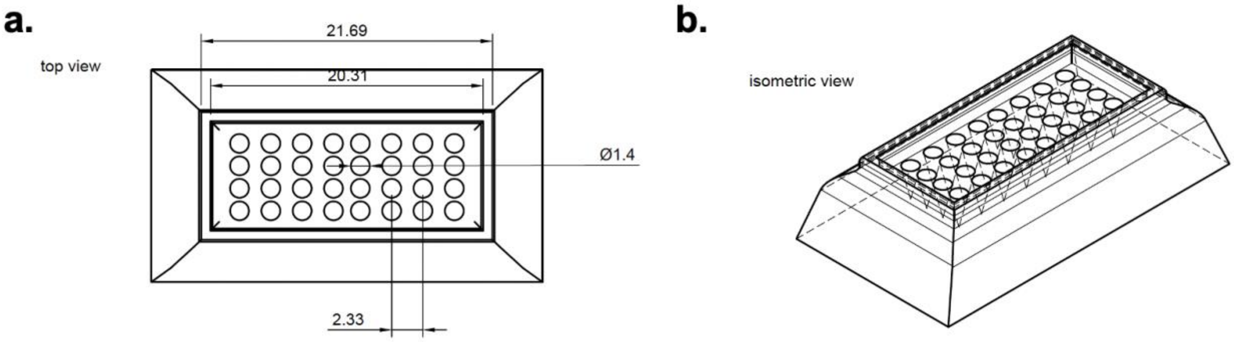


**Figure S2.** Schematic illustrations of PDMS negative mold.

**Table S1.** Summary of microneedle fabrication test conditions.

| Condition ID | Gelatin Conc. | Dehydration Time | Anthocyanin Conc. |
|---|---|---|---|
| 1 | 5% | 24 h | 0% |
| 2 | 10% | 24 h | 0% |
| 3 | 15% | 8 h | 0% |
| 4 | 15% | 16 h | 0% |
| 5 | 15% | 24 h | 0% |
| 6 | 20% | 24 h | 0% |
| 7 | 15% | 24 h | 0.1% |
| **8** | **15%** | **24 h** | **0.5%** |
| 9 | 15% | 24 h | 0.9% |

a.

b.

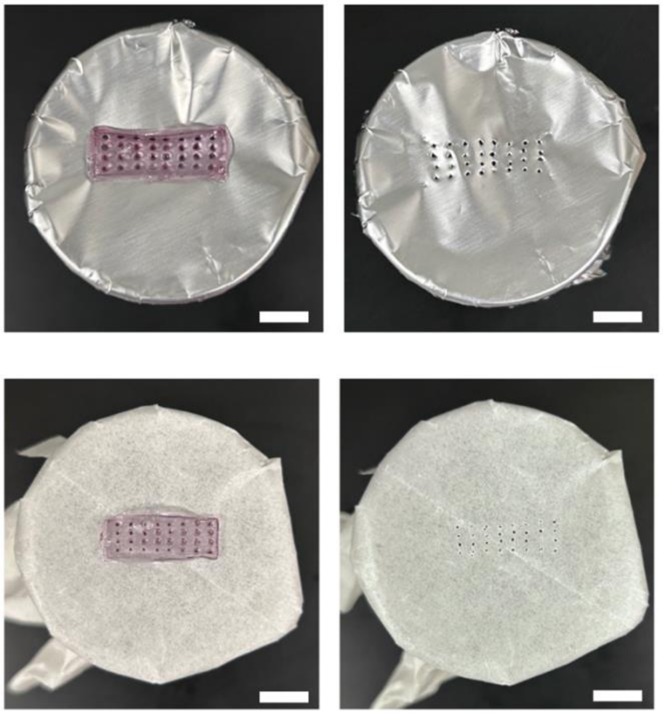

**Figure S3.** Penetration of alternative packaging materials. (a) Application and resultant penetration of tin foil. (a) Application and resultant penetration of wax paper. Scale bar depicts 10 mm.

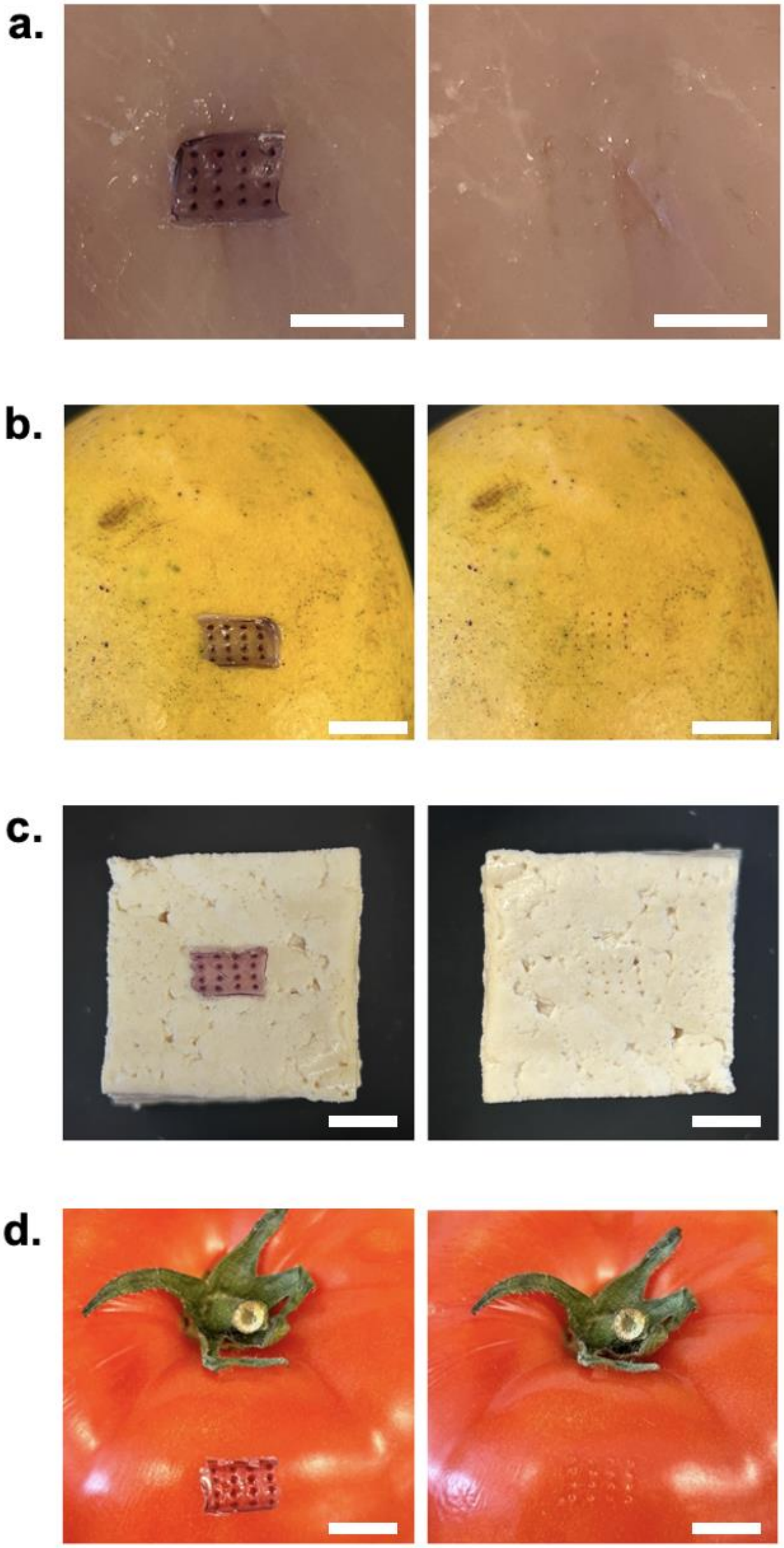


**Figure S4.** Microneedle application and resultant penetration on (a) chicken, (b) mango, (c) tofu, and (d) tomato. Scale bar depicts 10 mm.

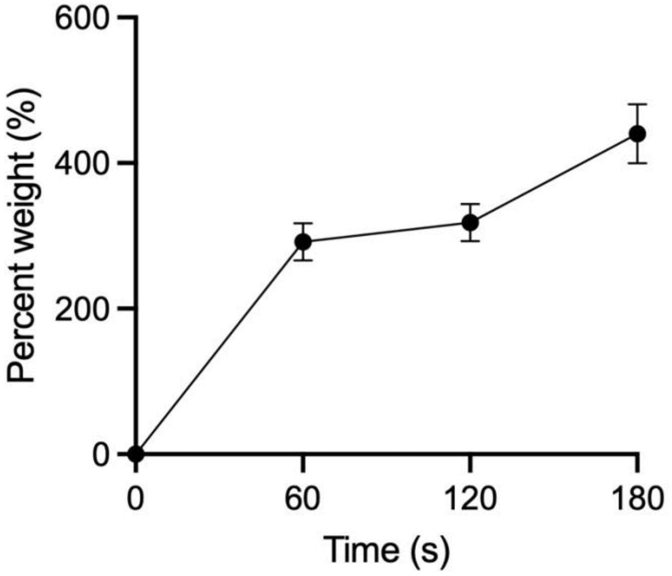


**Figure S5.** Dehydrated gelatin rehydration – defined by weight of water absorbed, across the first three minutes.

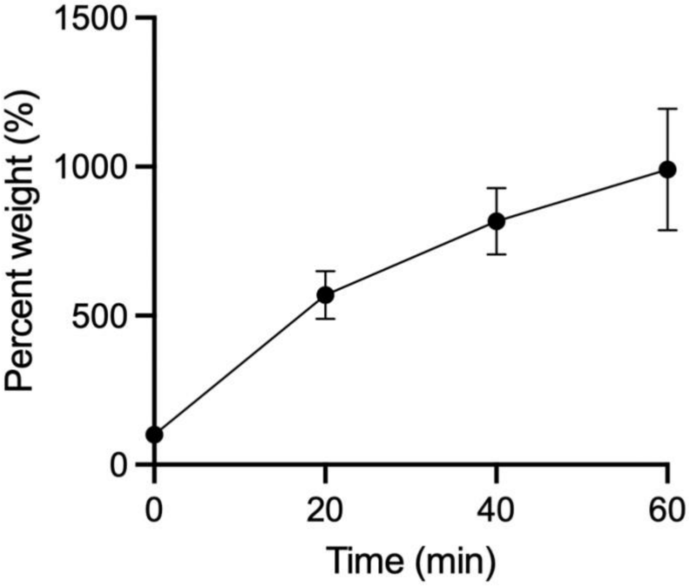


**Figure S6.** Dehydrated gelatin rehydration – defined by weight of water absorbed, across one hour.

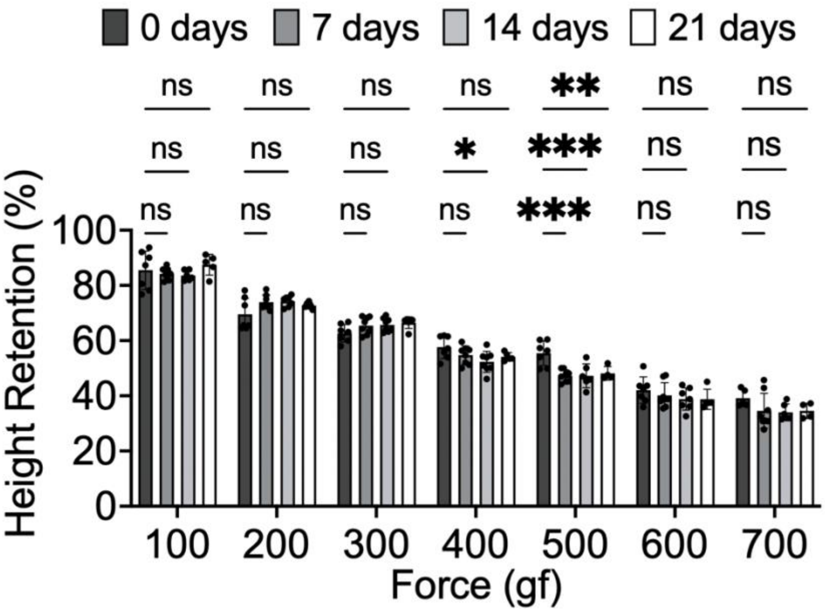


**Figure S7.** Mechanical integrity of 0.9% anthocyanin-embedded microneedles following storage at room temperature for varying lengths of time.

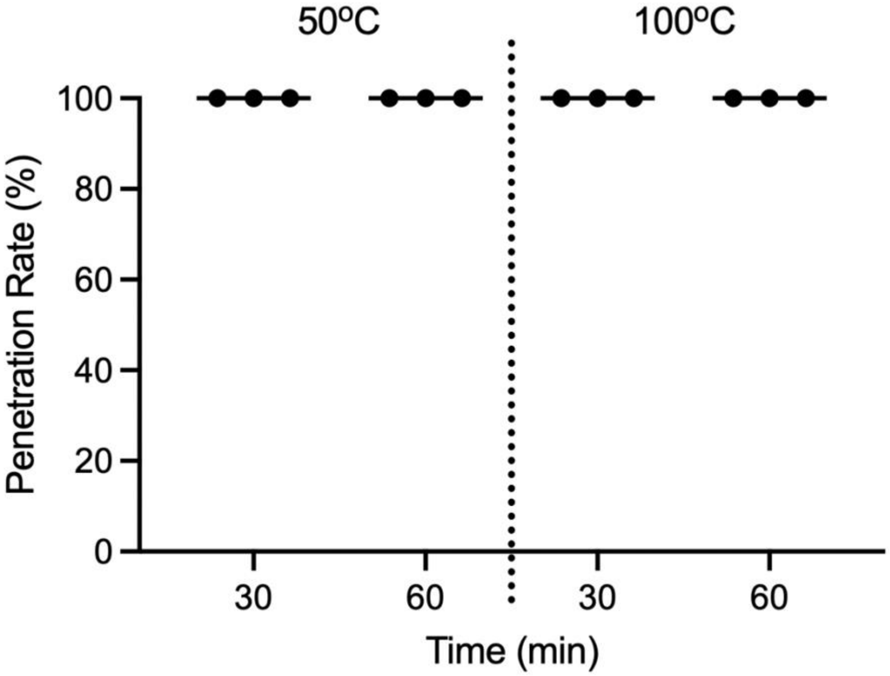


**Figure S8.** Dehydrated gelatin microneedle penetration rates following 30 min and 60 min exposure to high temperature conditions.

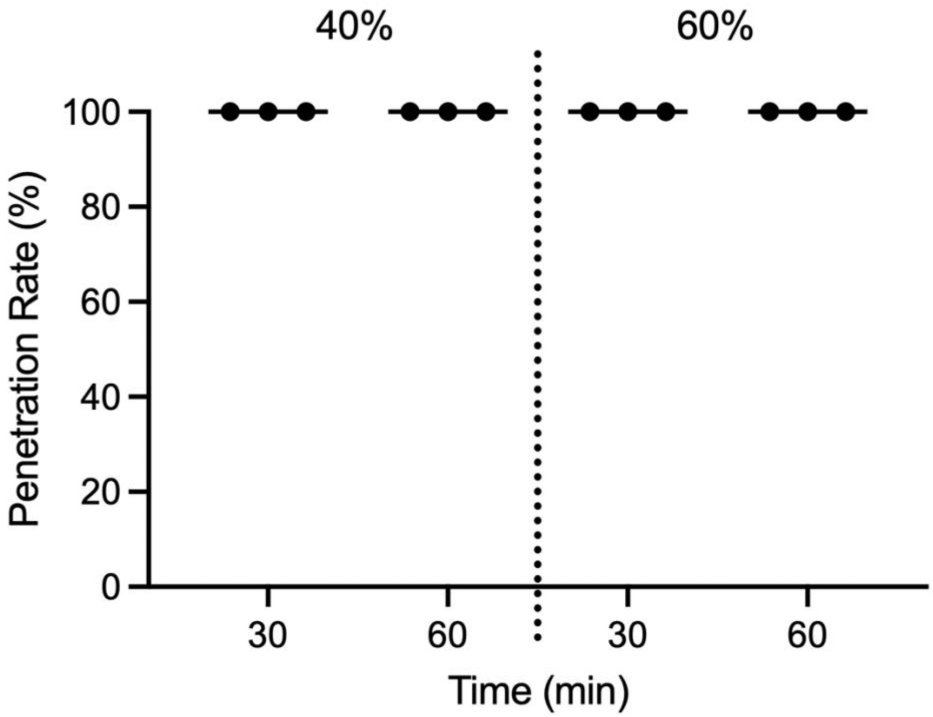


**Figure S9.** Dehydrated gelatin microneedle penetration rates following 30 min and 60 min exposure to varying humidity conditions.

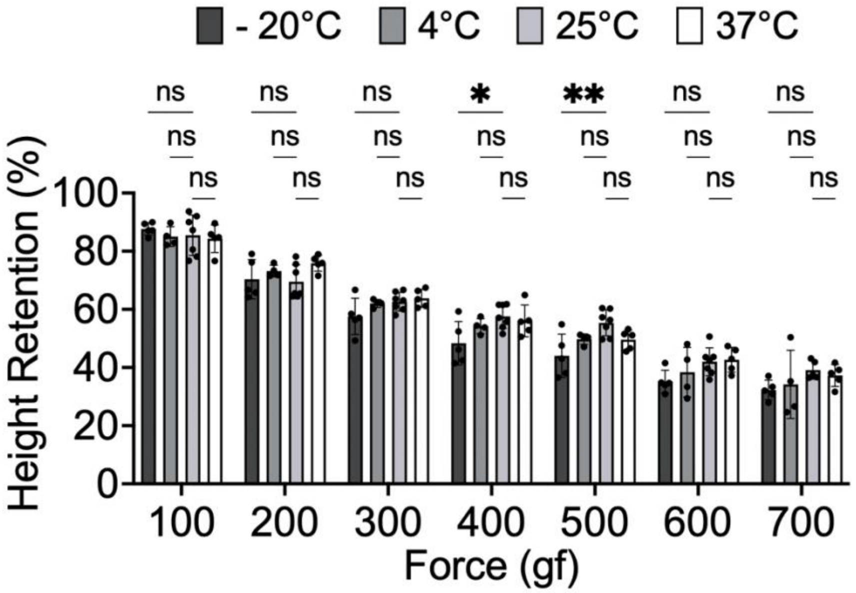


**Figure S10.** Mechanical integrity of 0.9% anthocyanin-embedded microneedles following exposure to diverse temperature conditions.

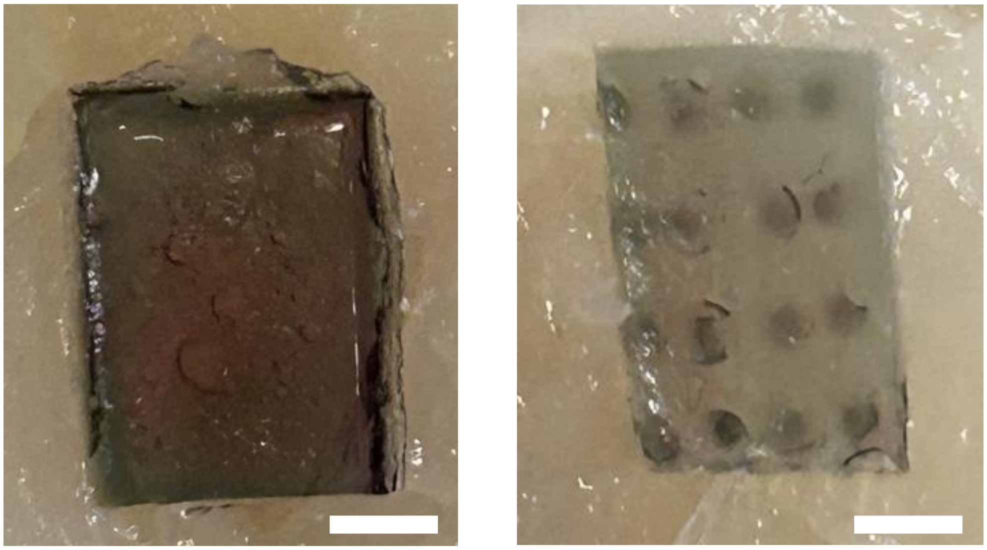

**Figure S11.** Colorimetric response of 0.5% anthocyanin-embedded, dehydrated flat patch (left) versus 0.5% anthocyanin-embedded, dehydrated microneedles after 45 minutes of insertion in an unpackaged, spoiled fish product. Scale bar depicts 5 mm.
